# Discovering Data Access and Use Requirements Using the Data Tags Suite (DATS) Model^1^

**DOI:** 10.1101/518571

**Authors:** George Alter, Alejandra Gonzalez-Beltran, Lucila Ohno-Machado, Philippe Rocca-Serra

## Abstract

This article presents elements in the Data Tags Suite (DATS) metadata schema describing data access, data use conditions, and consent information. DATS is a product of the bioCADDIE Project, which created a data discovery index for searching across all types of biomedical data. The “access and use” metadata items in DATS are designed from the perspective of a researcher who wants to find and re-use existing data. Data reuse is often controlled to protect the privacy of subjects and patients. We focus on the impact of data protection procedures on data users. However, these procedures are part of a larger environment around patient privacy protection, and this article puts DATS metadata into the context of the administrative, legal, and technical systems used to protect confidential data.

---

The vast amounts of data generated by researchers in many scientific disciplines hold potential discoveries extending beyond the work of those who created them. This is only possible if data can be discovered, accessed and amenable to reuse. The bioCADDIE Project (Ohno-Machado et al., 2017), which was funded by the NIH Big Data to Knowledge Program (BD2K) (Bourne et al., 2015) to create a way for researchers to search across all types of biomedical data, recognized that access conditions are an important part of data discovery. Researchers, when asked for cases for a data discovery index, emphasized their need for information about the conditions and methods for retrieving datasets of interest. Data accessibility continued to be a focus in work of the NIH Data Commons Pilot Phase Consortium that followed BD2K.

The possibility of data reuse is related to the use conditions for the data, and the latter has a direct impact on data privacy issues. An increasing number of research studies require data that may present risks to privacy if they are not protected. Human subjects may be exposed to re-identification from their genomes (Lippert et al., 2017), geographic locations (El Emam, Brown, & AbdelMalik, 2009), or clinical data (El Emam, Jonker, Arbuckle, & Malin, 2011), and pieces of information that may be innocuous in isolation can allow re-identification when combined, particularly when linked to other datasets. A wide range of procedures and technologies are being deployed to allow researchers to analyze these data while protecting rights of research subjects (Abowd & Lane, 2004; Sweeney, Crosas, & Bar-Sinai, 2015; Arellano, Dai, Wang, Jiang, & Ohno-Machado, 2018; Goroff, Polonetsky, & Tene, 2018). Researchers are aware that protecting confidential information imposes costs on them, and they want to know what to expect when it comes to data access and reuse conditions.

This article describes elements in the Data Tags Suite (DATS) metadata schema (Sansone et al., 2017, DATS 2018) designed to provide information ranging from data access to data use conditions and consent information. The “access and use” metadata items in DATS are designed from the perspective of a researcher who wants to find and re-use existing data. We focus on the authorization to use a dataset, and we do not attempt to describe the rules used to classify data as confidential or the characteristics of data that make them sensitive. We also do not examine technical aspects of authentication, i.e., confirming the identity of the researcher. Since DATS was created for use in a data discovery index, we emphasize the impact of data protection procedures on data users. However, these procedures are part of a larger environment around patient privacy protection, and this article puts DATS metadata into the context of the administrative, legal, and technical systems used to protect confidential data.

## Protecting Confidential Research Data

Data providers try to balance the benefits of facilitating access to existing research data with their obligation to protect information provided by research subjects (Alter & Gonzalez, 2018). Datasets vary in the level of risk that subjects can be re-identified and in the amount of harm that subjects would suffer if their confidential information became known. We also have a range of data protection measures that differ both in their effectiveness and in the costs that they impose on researchers. The most burdensome and costly types of data protection are normally for data that pose the greatest risks to subjects (Kaye & Hawkins, 2014; Sweeney, et al., 2015; Rubinstein & Hartzog, 2016).

Felix Ritchie (2005) proposed a framework for protecting confidential data that is known as the “Five Safes”.

- **Safe data:** Modify the data to reduce the risk of re-identification of subjects.
- **Safe projects:** Review and approve designs of proposed research projects.
- **Safe settings:** Isolate the data in a secure physical location or by applying secure remote access technologies.
- **Safe people:** Require legal agreements that commit researchers to protecting confidential information. Train researchers in best practices.
- Safe outputs: Review analyses and other products before releasing them to researchers. (Desai, Ritchie, & Whelpton, 2016)

These headings describe a toolkit from which data administrators select a combination of measures appropriate for the disclosure risks in a particular dataset (Broes, Lacombe, Verlinden, & Huys, 2018).

For example, a national sample of health interviews may be released after data masking procedures (“safe data”), such as “top coding” income into an open ended category to make very wealthy individuals less identifiable. In contrast, health histories of patients with a specific disease in a limited geographic area are much more difficult to de-identify. Patient records may be released for only approved types of research (“safe projects”) through a secure remote access system (“safe settings”) under a formal data use agreement (“safe people”). Additionally, some healthcare institutions only permit release of analyses performed on their data after review (“safe outputs”). The challenge for a data discovery index is capturing those aspects of the data protection that impose costs on prospective data users, and potentially displaying only datasets that users may be authorized to use. The user should be able to filter results according to his/her ability to conform to the authorization criteria.

## DATS Access Metadata

The bioCADDIE Project invited an international group of advisors to participate in an Accessibility Metadata for Datasets Working Group to recommend metadata describing how researchers gain access to data.^2^ This group identified three processes in accessing data for reuse: authorization, authentication, and access method.

### Authorization

Obtaining permission from the party that owns or is responsible for protecting the data is an important step. A range of checks can be done for credentialing a user, which may take milliseconds or stretch into weeks. Conditions for authorization are spelled out in data use agreements that are based on study consent forms, HIPAA authorization forms, and other documents. Some data created for public use may not require any kind of permission (open access), but confidential data are protected by formal authorization agreements, which are called ‘licenses’ in DATS (see below). For example, the U.S. Health Insurance Portability and Accountability Act (HIPAA) defines two standards for disclosing protected health information: “de-identification” and development of ‘limited data sets’ (U.S. Department of Health and Human Services, 2000). ‘Limited data sets’ are only available under a data use agreement, because they contain information that increases the risk of re-identifying individuals.^3^ The most restrictive authorization procedures are designed to limit data access to “safe people” (controlled access) who will respect the rights of research subjects and patients. Higher levels of security also come with a price. Obtaining institutional signatures on legal agreements is burdensome and reduces re-use of data (Joly, Dyke, Knoppers, & Pastinen, 2016; Kaye & Hawkins, 2014). For electronic health record data, researchers are typically not the signatories of Data Use Agreements (DUAs): this is usually reserved for institutions. Dyke et al. (2016) propose registration with self-declaration of qualifications, purpose, and commitments as a level of protection between open access and authorization under formal agreements. The Working Group identified six common types of authorization.

| Authorization Types |  |
| --- | --- |
| **None** | Not covered by a data use agreement. |
| **“Click through” online license** | Users must agree to an online agreement without providing additional identification. |
| **Registration** | Users must register before access is allowed and agree to conditions of use. Registration information may be verified. |
| **Data use agreement signed by an individual** | An agreement signed by the investigator is required. Data use agreements may require additional information, such as a research plan and an Institutional Review Board (IRB) review. (See discussion of Licenses below.) |
| **Data use agreement signed by an institution** | An agreement signed by the investigator's institution is required. Data use agreements may require additional information, such as a research plan and an IRB review. (See discussion of Licenses below.) |

Authorization procedures have implications for the accountability of the data user and timelines for accessing the data. For example, if researchers want to remain anonymous they can only access datasets labeled with Authorization Types “None” or “Click through.” If they are not affiliated with an institution, they are not eligible for data use agreements covering many types of biomedical data. Conditions in the DUA impose additional hurdles before the researcher will be able to use the data. Approval by an institutional review board (IRB) is often required, and the researcher may need to show that the purpose of the research is consistent with the consent forms signed by participants in the study. Authorization is thus complicated, and a researcher may be allowed to use a particular dataset for one purpose (e.g., cancer research) but not another (e.g., a study on ancestry). (See the discussion of Licenses below.)

A data repository may use one or more of these authorization types. The Inter-university Consortium for Political and Social Research (2018), which is the oldest repository of social science data in the U.S., has examples of all five authorization types among the studies in its collection.

### Authentication

When data are accessed online, many data repositories require some kind of login process to identify the user. Even when the data are not covered by a license, the user may need to create a username and password for access (i.e., registered access). Access to confidential data may require multi-factor authentication controls involving a second type of identification, such as a telephone number or dedicated IP address. Researchers who plan to automate harvesting of data from multiple sources are especially interested in authentication procedures. Three types of authentication were listed by the Accessibility Metadata for Datasets Working Group.

| Authentication types |  |
| --- | --- |
| None | No authentication required. |
| Simple login | Single factor login or the use of an authentication key or registered IP address is required. |
| Multi-factor login | Multiple factor login using a combination of IP address, password protection, authentication key, or other forms of authentication. |

### Access Method

Data repositories may protect confidential data by only allowing access in a physical or virtual “safe setting”. Researchers who want to use highly sensitive data may need to travel to a secure “enclave”, such as the Census Bureau’s Research Data Centers (U.S. Census Bureau, 2018) and the Veterans Health Administration VINCI system (U.S. Department of Veterans Affairs, 2018), or submit program code to be executed by the data repository (“remote service”) (Abowd & Lane, 2004). A growing number of data providers allow researchers “remote access” to computers in a secure data center, and this is the model selected by the NIH-funded AllofUs Research Program (National Institutes of Health, 2017, p. 55). Researchers working in these “virtual data enclaves” see a standard operating system, as they would on their local computer, but they cannot download data to their local machine (Data Sharing for Demographic Research, 2018; Research Data Assistance Center, 2016). They are a virtual machine launched from their local computer but actually operating on the remote system. An example of this is Vivli (Bierer, Li, Barnes, & Sim, 2016), a new platform for sharing clinical research data, connects a data repository to a secure cloud-based workspace. The “Data Commons” being developed by NIH will also rely, at least in part, on a remote access system. When researchers are required to perform all of their analyses on a computer controlled by the data provider, the provider also has the option of examining and approving results before sending them to the researcher (“safe outputs”).

| Access Method |  |
| --- | --- |
| Download | The data are available for download. A license may be required. |
| API | Interaction with the data may be automated via defined communication protocols, i.e., Applications Programming Interfaces (APIs). |
| Remote access | Users may access the data in a secure remote environment (“virtual data enclave”). Individual-level data may not be downloaded, only approved results. |
| Remote service | A user may submit program code or the script for a software package to be executed in a secure data center. The remote site returns outputs. It may perform a review before releasing the results. |
| Enclave | Access is provided to approved users within a secure facility without remote access. Results may remain at the enclave or be released after review. |

## DATS Use Metadata

A variety of terms are used to refer to legal agreements between data providers and users, including “data use agreement,” “data access agreement,” “material transfer agreement,” and “non-disclosure agreement.” In the “open data” world these agreements are called “licenses.” For example, Creative Commons, which is best known for its copyright licenses, also provides licenses for data (Creative Commons, 2018). However, “license” usually implies access to a commercial product (e.g., software) or intellectual property (e.g., an invention), and we prefer “data use agreement” (DUA) for the agreements between data providers and users, especially for confidential data.

Data use agreements are often lengthy agreements that inherit conditions from a number of earlier documents involving several different parties (see Figure 1). DUAs include provisions describing allowed uses, limitations, and requirements: what analyses can be conducted, how long the data may be used, and the ways in which it must be returned, destroyed or discarded after use. DUAs typically require data users to obtain Institutional Review Board (IRB) approval from their home institutions, which may impose additional conditions on the data user’s research plan. The bioCADDIE project did not attempt to create a comprehensive ontology of conditions found in data use agreements, but DATS has been designed to take advantage of current efforts to use relevant ontologies when available. To understand the metadata needed to describe an agreement for re-using data, we outline how these agreements are created and administered. We focus here on the most complex case: agreements for data with some risk of harm to research subjects or patients.

**Figure 1.**
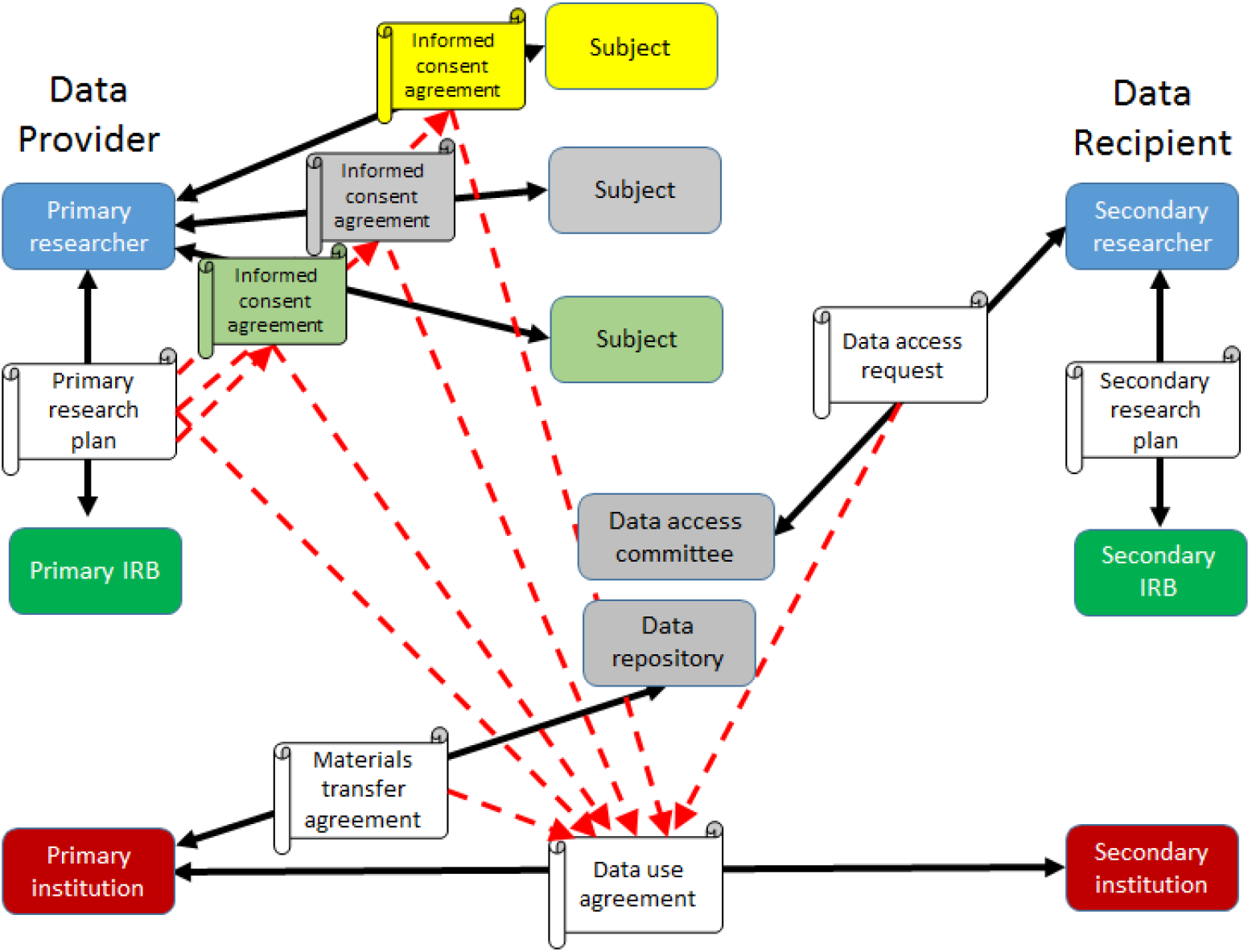
The network of agreements from data collection to data sharing.

Most academic research involving human subjects requires prior approval of a research plan by an ethical review committee or an IRB. In the U.S., federal regulations mandate IRB approval for all research sponsored by NIH, NSF, and some other agencies (U.S. Department of Health and Human Services, 2009), and most universities and other organizations require IRB review for all research involving human subjects. IRBs are responsible for protecting human subjects from the risks posed by research, which they do by approving and monitoring compliance with research plans. The research plan will include a description of documents and procedures for obtaining informed consent from research subjects involved in the study. The terms of the informed consent apply to all future research, and an IRB should also review plans for sharing data resulting from the study. Researchers who analyze confidential data from a data repository are also expected to obtain approval from an IRB at their institution. However, IRBs do not make agreements with researchers at other institutions to share confidential data.

Transactions involving confidential research data are conducted by officials who are authorized to make agreements for the institution, such as a research administration officer. Most universities assert that research data belong to the institution not to the researcher. If the research was sponsored by a funding agency, the ‘grantee’ is the university, and universities see ownership of data resulting from external funding as part of their obligation to assure compliance with the terms of the grant or contract. Since confidential data also pose a risk to the reputation of the institution and possibly legal liability, universities are especially motivated to monitor the agreements surrounding them. This also applies to the institutions of researchers who request confidential data, and many universities will not allow faculty or staff to sign DUAs. Since data providers want the recipient university to be responsible for the management of confidential data, data use agreements are typically signed by university officials on both sides.

Data use agreements often include a variety of conditions that were not explicitly included in the Informed Consent agreement. For example, some agreements include detailed requirements about data storage and computer systems. Agreements may require data to be stored offline and isolated from the Internet or be encrypted. Data recipients are often required to inform the data provider about any publications resulting from their secondary research. In some cases, the data provider insists on reviewing articles before they are submitted for publication or public presentation. Most data providers require a research plan describing how the data will be used, which may be reviewed by a panel of experts. For instance, researchers who ask for data from NIH’s Database of Genotypes and Phenotypes (dbGaP) repository must submit a Data Access Request for review by a Data Access Committee (Paltoo et al., 2014; Shabani, Dove, Murtagh, Knoppers, & Borry, 2017). The Data Access Request becomes part of the Data Use Agreement and limits the recipient to the approved analyses.

‘Dynamic consent’ is a rapidly developing practice with important implications for data access (Budin-Ljosne et al., 2017). Until recently, informed consent agreements were static documents signed by a research subject when data were collected. There is a strong movement to give research subjects ongoing control over the use of their data (Genetic Alliance, 2018). Subjects may be able to withdraw consent at any time, and several new technologies allow them to choose which research projects can use their data (Kim et al., 2017; Wilbanks & Friend, 2016). Dynamic consent conforms to the spirit of the European Union’s General Data Protection Regulation (GDPR), but the GDPR exempts scientific research from rules giving subjects control of their data (Chassang, 2017; European Union, 2016). This is for research considered of “substantial public interest”, in which case consent is not required. If the research is not considered in the public interest, there are more demanding requirements entailing true anonymization of the data or consent (Rumbold and Piersioneck 2017).

Private companies now hold enormous quantities of confidential data about their customers, which are sometimes available to academic researchers under data use agreements. Kanous and Brock (2015) found that agreements used by private data providers were often poorly designed. These agreements were usually derived from non-disclosure or confidentiality agreements that were designed to protect the business secrets of the data provider. Consequently, they are often vague about the nature of the data and the uses permitted to the researcher. Kanous and Brock (2015) also found that some agreements include conditions asserting the data provider’s right to ‘derivative’ works, which might be interpreted to include analyses and publications. The Yale Open Data Access (YODA) Project was developed to provide independent scientific review of requests to use data created in the private sector (Krumholz & Waldstreicher, 2016).

The Accessibility Metadata for Datasets Working Group decided not to attempt to characterize the conditions included in a data use agreements. Creating an ontology of use conditions was deemed beyond the resources of the group. Fortunately, since the Working Group finished its report, a number of efforts have moved in the direction of ontologies describing the conditions in data use agreements, which will be detailed below. To accommodate this type of information, the most recent version of DATS was extended with ‘dataUseCondition’ and ‘consentInformation’ schemas for referencing dedicated ontologies in anticipation of their future implementation.

### Conditions in Data Use Agreements

Data providers use agreements to assure that data users behave in ways that protect confidential information and respect consent agreements with research subjects. For example, a common requirement is that researchers will not attempt to re-identify subjects (“safe people”). Data use agreements used by the Inter-university Consortium for Political and Social Research (ICPSR) include lists of statistics that should not be published, such as a cell in a cross-tabulation table describing only one person, because of re-identification risks. Some agreements require data recipients to submit papers and presentations for review before publication. Standardization of data use conditions would make it easier to automate management and compliance with data use agreements, but the diversity and specificity of legacy agreements makes classification very difficult. Fortunately, several projects are working on this problem.

“Automatable Discovery and Access Matrix” (ADA-M) is an ambitious project of the Global Alliance for Genomics and Health (GA4GH) and the International Rare Diseases Research Consortium (IRDiRC) to standardize metadata about data access (Woolley, 2017; Woolley et al., 2018). ADA-M divides conditions into “permissions” and “terms,” which are arranged under specified concepts. Additional information about permissions and terms can be provided with child fields, and free-text fields are used to capture details and for human readability.

The Data Use Ontology (DUO), which is also a project of GA4GH, is formalizing a controlled vocabulary used by dbGAP for conditions in data use agreements for genomic data (Dyke, Philippakis, et al., 2016). DUO is based on the NIH Standard Data Use Limitation (DUL) codes (National Institutes of Health, 2015).

Data in dbGAP studies are arranged into ‘consent groups’ that share consent agreements and other use conditions. DULs are used to summarize these conditions, although consent groups are often subject to additional conditions not described by DULs. DUL codes are composites combining several types of conditions. For example, General Research Use (GRU) the broadest DUL code allows studies of statistical methods and population structure or ancestral origin, but the Health/Medical/Biomedical (HMB) code excludes those studies. Like the DUL codes, DUO allows a primary category (e.g., GRU) to be modified by a secondary category (e.g., NMDS no general methods research). DUO has been adopted by the European Genome-Phenome Archive (European Genome-Phenome Archive, 2018).

The Informed Consent Ontology (ICO) is being developed to represent logically terms and relations in informed consent agreements (Lin et al., 2014; Manion et al., 2014). As the process of obtaining informed consent moves from paper to online systems, it becomes possible to offer individual subjects a wider range of choices about the future use of their data.

The Open Digital Rights Language (ODRL), a recommendation by the World Wide Web Consortium (W3C) (Ianella et al 2018), provides a rich language to express statements about the usage of content and services. It allows to provide a standard description model and format representing permission, prohibition, and obligation statements. ODRL is recommended in the Data Catalog Vocabulary (Gonzalez-Beltran et al 2018).

Other relevant vocabularies, which also follow a granular approach, are the Open Data Rights Statement Vocabulary (Dodds 2013) and the Agreements ontology (AGR-O) (Car 2018).

These vocabularies provide more granularity than DUO and ICO. However, representing Data Use Agreements or Data Use Limitations in a granular way distinguishing permissions, prohibitions and obligations is a very challenging task, which becomes intractable in many cases, as the original documents have not considered such detailed representation and there is ambiguity on choosing relevant terms.

Thus, for cases where it is not feasible to distinguish between permissions, prohibitions and obligations, DATS recommends the use of DUO and ICO. However, if an expression relying on ODRL, or ODRS, or the Agreements ontology can be used, DATS support pointing to such expression.

The DATS ‘ConsentInformation’ (consent_info_schema.json)^4^ schema has been designed in a flexible way to capture conditions limiting the use of a dataset that may not be included in the data use agreement. First, as we noted above, the data use agreement implicitly inherits conditions from all of the previous agreements and approvals covering the data. In particular, the informed consent agreement may include requirements not listed explicitly in the data use agreement. For this reason, the ‘consentInformation’ schema includes an ‘incorporatedIn’ property that points to the license that it modifies (see Figure 2).

**Figure 2:**
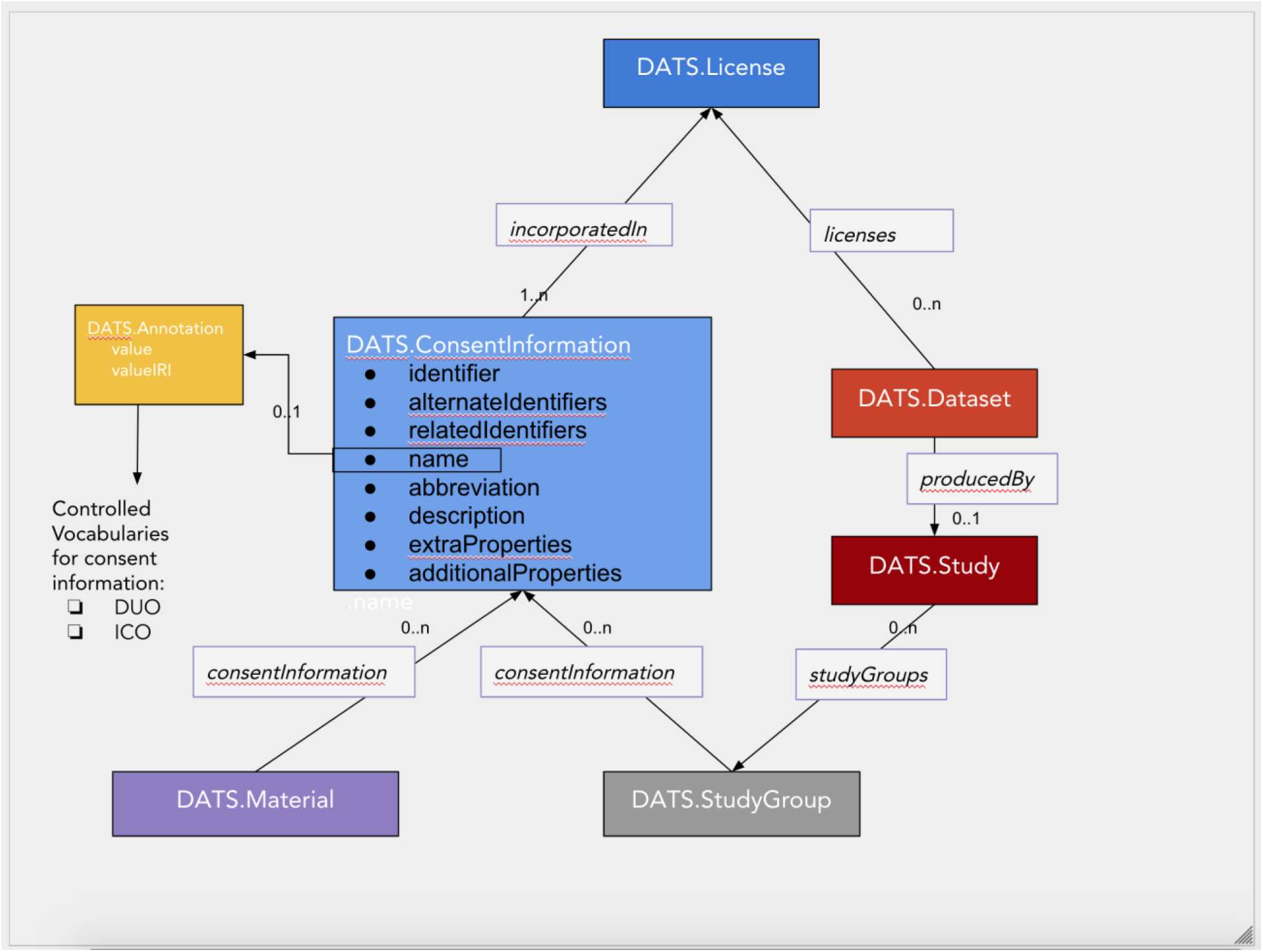
a graphical representation of relevant constructs allowing consent, license and terms of use information to be made available as information payload in DATS messages. The new *ConsentInformation* schema allows for annotation (semantic markup) with resources such as the Data Use Ontology (produced by the Global Alliance for Genomic Health) or the Information Consent Ontology.

Second, a research study may include data from subjects who signed different consent agreements. An important example of this regularly occurs in studies that collect genomic data from patients with a specific disease. Some subjects provide consent only to research about their disease, while other subjects allow their data to be used for any type of research. Restrictions of this kind are important to researchers who are searching for data as well as collecting data for reuse (e.g., when creating synthetic cohorts). DATS may be used to describe a specimen or data derived from a tissue sample of a specific individual. To cover these cases the ‘consentInformation’ property may be included in the DATS ‘material’ entity.

The following table shows the information in dbGAP for consent groups in the Massachusetts General Hospital (MGH) Atrial Fibrillation Study. The first group (Health/Medical/Biomedical) gave their consent for any type of health, medical, or biomedical research with the exception of studies about the origins or ancestry of individuals or groups. The second group (Disease-Specific) only consented to future research on atrial fibrillation, the focus of the original study. Both consent groups require IRB approval from the recipient’s institution. In these cases, DATS can describe multiple “study groups” with different consent conditions and other attributes. In DATS, we would represent each consent group as a StudyGroup with different ‘consentInformation’.

| Consent group | Consent Information |
| --- | --- |
| Health/Medical/Biomedical (IRB) | Use of this data is limited to health/medical/biomedical purposes, does not include the study of population origins or ancestry. Requestor must provide documentation of local IRB approval. Use of the MGH AF Study data deposited in dbGaP is restricted to research on associations between phenotypes and genotypes. MGH AF Study data may not be used to investigate individual participant genotypes, individual pedigree structures, perceptions of racial/ethnic identity, non-maternity/paternity, and of variables that could be considered as stigmatizing an individual or group. All research must be related to the etiology and prevention of morbidity and mortality of the U.S. population consistent with the demographic distribution in the MGH AF Study. Data users will be required to obtain IRB approval for their projects from their respective institutions (please note that only full or expedited approvals will be accepted). |
| Disease-Specific (Atrial Fibrillation, IRB, RD) | Use of the data must be related to Atrial Fibrillation and related disorders. Requestor must provide documentation of local IRB approval. Data use is limited to research related to atrial fibrillation and cardiovascular disease. |
| Source: NHLBI TOPMed: Massachusetts General Hospital (MGH) Atrial Fibrillation Study<br>dbGaP Study Accession: phs001062.v1.p1. https://www.ncbi.nlm.nih.gov/projects/gap/cgi-bin/study.cgi?study_id=phs001062.v3.p2 |  |

Third, we expect that greater standardization and automation of informed consent agreements will lead to the use of ontologies describing the conditions within these agreements. DATS has a standard way of referring to external ontologies, which can be used for the standards being developed by projects like ADA-M, DUO, and ICO. By including these conditions in DATS, we allow them to be used for discovery and for filtering search results. Standardization will make it easier to provide this important information to researchers.

To record aspects of dynamic consent, the ‘ConsentInformation’ schema also has a property for ‘temporalCoverage’, allowing to indicate the periods of time when the consent is valid.

## Discussion

There is an inherent tension between the growing importance of research that combines data from multiple sources and the increasing demand for data that cannot be de-identified. Researchers cannot plan their work unless they know how access will be provided and how long it will take to obtain the necessary permissions. The access metadata objects in the DATS metadata standard differ from other approaches in their focus on the experience of researchers who need to find and intend to re-use existing data. DATS access metadata does not have the level of detail found in metadata standards designed for managing data resources, like ADA-M (Woolley, et al., 2018), Fast Healthcare Interoperability Resources (FHIR) (Health Level Seven International (HL7), 2018), or eXtensible Access Control Markup Language (XACML) (OASIS TC, 2018), but references to other metadata standards can be embedded in DATS. As these and other standards and ontologies develop, data discovery applications will be able to benefit from them through DATS.

Based on the experience garnered through work on the DataMed prototype and the Data Commons Pilot Project Consortium, we are convinced that dividing the access process into three steps (authorization, authentication, and type of access) is a useful and original contribution of DATS. New ways of implementing each of these steps are still emerging. Most discussions of data access distinguish between ‘open’ and ‘restricted’ data, but ‘restricted’ data are distributed in a growing number of different ways. From a researcher’s point of view data that can be downloaded are very different from data that are only accessible on a remote virtual machine. As the bioCADDIE Project has drawn to a close, we document our experience and encourage other organizations to take responsibility for supporting and updating the controlled vocabularies identified by the Accessibility Metadata for Datasets Working Group.

Capturing metadata about the conditions affecting data use will be a time consuming process until standard ways of describing informed consent and data use agreements become part of automated systems for creating and managing research data. There is little standardization in these agreements today, and extracting and classifying the conditions included in legacy agreements is a very complex task. When agreements have been described in standards like ICO, DUO, and ADA-M, they will be searchable and discoverable in DATS. We expect that the benefits of automating these agreements will be great, but it will take time to be realized. Since the technology for electronic health records is developing very quickly, the automation of consent for research use of patient records and tissue samples in FHIR or other standards may be close.

The most difficult problem is obtaining the cooperation of data providers in describing their access and licensing procedures. This does not mean that all data providers must expose metadata about their holdings in DATS. The bioCADDIE Project has demonstrated the flexibility of DATS by mapping and ingesting metadata from more than 70 data repositories into DataMed (Chen et al., 2018). However, the capabilities of data repositories vary widely. Major data repositories (e.g., dbGAP, Protein Data Bank, ICPSR) have established metadata standards as well as the material and human resources to adapt to new requirements. Other data repositories operate with minimal staff and under precarious funding, even if they serve important scientific communities. We see a great need for NIH and other funding agencies to adopt standards for data repositories, such as the CoreTrustSeal (CoreTrustSeal, 2018), and develop new funding mechanisms designed to provide sustainable support for data curation, dissemination, and preservation. As funding agencies put increased emphasis on FAIR (Findability, Accessibility, Interoperability, Reusability) principles (Wilkinson, et al., 2016), access and data use conditions should become findable as well.

## Footnotes

2 We are very grateful to the members of bioCADDIE Project Working Group 7 “Accessibility Metadata for Datasets”: George Alter (chair, University of Michigan), Damon Davis (HealthData.gov), Alex Kanous (University of Michigan), Hyeoneui Kim (University of California San Diego), Jared Lyle (University of Michigan), Frank Manion (University of Michigan), Reagan Moore (University of North Carolina), Mark Phillips (McGill University), Kendall Roark (Purdue University), Jessica Scott (GlaxoSmithKline), Anne-Marie Tasse (McGill University).

3 Several authors have shown that individuals can be re-identified in “de-identified”. See, for example, (El Emam, et al., 2011).

4 https://github.com/datatagsuite/schema/blob/master/consent_info_schema.json

## Acknowledgment

This work was supported by the bioCADDIE Project, funded by NIH grant 1U24AI117966-01. We are very grateful to Elaine Brock, Alex Kanous, and Melanie Courtot for valuable comments on a previous draft of this article.

